# pepFunk, a tool for peptide-centric functional analysis in metaproteomic human gut microbiome studies

**DOI:** 10.1101/854976

**Authors:** Caitlin M.A. Simopoulos, Zhibin Ning, Xu Zhang, Leyuan Li, Krystal Walker, Mathieu Lavallée-Adam, Daniel Figeys

## Abstract

Enzymatic digestion of proteins before mass spectrometry analysis is a key process in metaproteomic workflows. Canonical metaproteomic data processing pipelines typically involve matching spectra produced by the mass spectrometer to a theoretical spectra database, followed by matching the identified peptides back to parent proteins. However, the nature of enzymatic digestion produces peptides that can be found in multiple proteins due to conservation or chance, presenting difficulties with protein and functional assignment. To combat this challenge, we developed a peptide-centric metaproteomic workflow focused on the analysis of human gut microbiome samples. Our workflow includes a curated peptide database annotated with KEGG terms and a pathway enrichment analysis adapted for peptide level data. Analysis using our peptide-centric workflow is fast and identifies more enriched KEGG pathways than protein-centric analysis. Our workflow is open source and available as a web application or source code to be run locally.

## Introduction

Metaproteomics, the study of proteins from an environmental sample, is used to examine the dynamics and composition of microbial communities in complex environments including human and animal microbiomes (Moon et al., 2018; Cheng et al., 2018), soil (Starke et al., 2019) and water samples (Mikan et al., 2019). Understanding the microbial dynamics and functionality of the human gut microbiome is particularly of interest due to its association with human disease as observed in immune-system associated diseases such as inflammatory bowel disease (Morgan et al., 2012; Zhang et al., 2018b), asthma (Arrieta et al., 2015), and multiple sclerosis (Jangi et al., 2016), metabolic disorders such as obesity and type II diabetes (Sonnenburg and Bäckhed, 2016), and cardiovascular disease (Tang et al., 2017). Studies have also demonstrated that the presence of the “gut-brain” axis can mean that gut microbes are capable of influencing, or are at least linked to, one’s mental health, with evidence even suggesting that modulation of microbiota can have therapeutic effects in anxiety and depression (Dash et al., 2015).

Although the term “proteomics” implies that proteomic data inherently consist of protein level information, an important step in most proteomic workflows is to enzymatically digest extracted proteins into smaller peptide fragments before mass spectrometry (MS) sequencing (Hettich et al., 2013). To facilitate the analysis, peptides are then separated and analyzed, often by liquid chromatography (LC) coupled with tandem MS. Raw spectra produced by MS/MS are computationally matched with predicted spectra of peptide sequences by database search. These matched peptides are then assigned to proteins. However, due to the nature of enzymatic digestion, the same peptide sequence can belong to multiple proteins, and it is difficult to determine the correct parent protein of these redundant peptides (Nesvizhskii and Aebersold, 2005). Nesvizhskii and Aebersold (2005) deemed this challenge the Protein Inference Problem, which is further exacerbated in metaproteomics experiments due to the presence of multiple microbial strains and species that can include additional redundant peptides due to protein sequence conservation. Nonetheless, computational workflows for proteomic research typically use proteins identified from peptide sequences for quantitative and functional enrichment studies although redundant peptides can impede accurate and confident identification of proteins from MS/MS data.

Ning et al. (2016) describe the uncertainty of peptide-to-protein assignment as “information degeneration”. This information loss stems from the methods that researchers have previously used to mitigate the ambiguity of peptide-to-protein assignment. For example, the Occam’s razor principle relies on discarding proteins without unique peptides, and often can only identify protein groups (Serang and Noble, 2012). Alternatively, Muth et al. (2015) have introduced the concept of a “meta-protein”, where proteins are grouped by amino acid sequence or shared peptides. Although methods have been introduced to combat the Protein Inference Problem, methods for a peptide-centric metaproteomic workflow have also been implemented to circumvent information loss. Notably, UniPept is a Gene Ontology (GO) term-focused functional analysis tool that was released as both a web application and local software tool (Gurdeep Singh et al., 2019). UniPept uses a large GO term functional database consisting of tryptic peptides of proteins found in the UniProt Knowledgebase. However, GO term annotations are organized in a directed acyclic graph with semantic relationships between terms, causing challenges for functional enrichment analyses such as unclear hierarchies and dependencies (Gaudet and Dessimoz, 2016). To manage this challenge, Riffle et al. (2018) created MetaGOmics, a peptide-centric GO term based enrichment tool that creates directly acyclic graphs for GO terms associated to identified peptides. Despite the computational progress these previously mentioned tools have made for peptide-centric metaproteomic workflows, the tools all use complex GO term annotation and are not specifically created for gut microbiome studies.

In this work, we introduce a novel peptide-centric workflow for metaproteomics data for gut microbiome experiments. To facilitate this work, we created a Kyoto Encyclopedia of Genes and Genomes (KEGG)-peptide functional database, a functional enrichment workflow and an interactive web application companion tool for gut microbiome metaproteomic studies. We created a peptide-function database consisting of *in silico* digested peptides from the Integrated human gut microbial Gene Catalog (IGC) database. We reduced the size of our peptide database by focusing on the most empirically identified peptides in raw MS/MS data for improved computational speed. We used annotation from UniProt Reference Clusters (UniRef90) sequence clusters to functionally annotate the gut microbiome peptide database with KEGG terms. We created a peptide-centric functional enrichment workflow by adapting gene set variation analysis (GSVA) for peptide-level data (Hänzelmann et al., 2013). We found that the results from our peptide-centric workflow correlated with results from a protein-centric workflow suggesting that the peptide workflow is suitable and comparable to a more canonical approach to metaproteomics data analysis. Additionally, our peptide-centric workflow was able to identify more enriched KEGG pathways than when using protein-level data. Finally, we packaged our peptide-centric data analysis pipeline into a user-friendly web application intended to be used as a companion tool to MetaLab and iMetaLab (Cheng et al., 2017; Liao et al., 2018), and released the source code to allow local data analysis for experienced computational users.

## Methods

### KEGG core peptide database construction

We used the IGC protein dataset (https://db.cngb.org/microbiome/genecatalog/genecatalog_human/; Li et al., 2014) to create our KEGG-peptide functional database. We first annotated the IGC protein dataset by searching for sequence identity in UniRef90 sequence clusters (Suzek et al., 2014). Sequence alignment was computed using Diamond blastp (Buchfink et al., 2015) and command line options `--sensitive -e 0.1 --top 5 -f 6 qseqid qlen sseqid slen evalye length mident`, where `--sensitive` gave us a search with a higher sensitivity, `-e 0.1` allowed for a maximum E value of 0.1, `--top 5` produced a list of the top five hits, and `-f` let us customize the output file. We considered a single protein match to a cluster sequence in UniRef90 as the smallest E value representing the best match for functional identification. Notably, 99.5% of protein matches have E values <0.0001. Each protein in the IGC dataset was then annotated with KEGG terms using the annotation associated to UniRef90 protein matches. We completed an *in silico* trypsin digestion of the IGC protein dataset using a Python script (https://github.com/northomics/bin/blob/master/trypsin.py) that considered digestion at lysine and arginine except if followed by a proline, and up to two missed enzymatic cleavages. Computed peptides inherited the KEGG functional annotation of parent proteins. In the case of redundant peptides, or peptide sequences that are found in multiple proteins, the union of all identified KEGG annotations for all parent proteins were considered (Figure 1a). In other words, we did not discard any putative functional annotation in redundant peptides and instead multiple KEGG annotations were accounted for in functional enrichment analysis by intensity weighting (Figure 1b).

**Figure 1:**
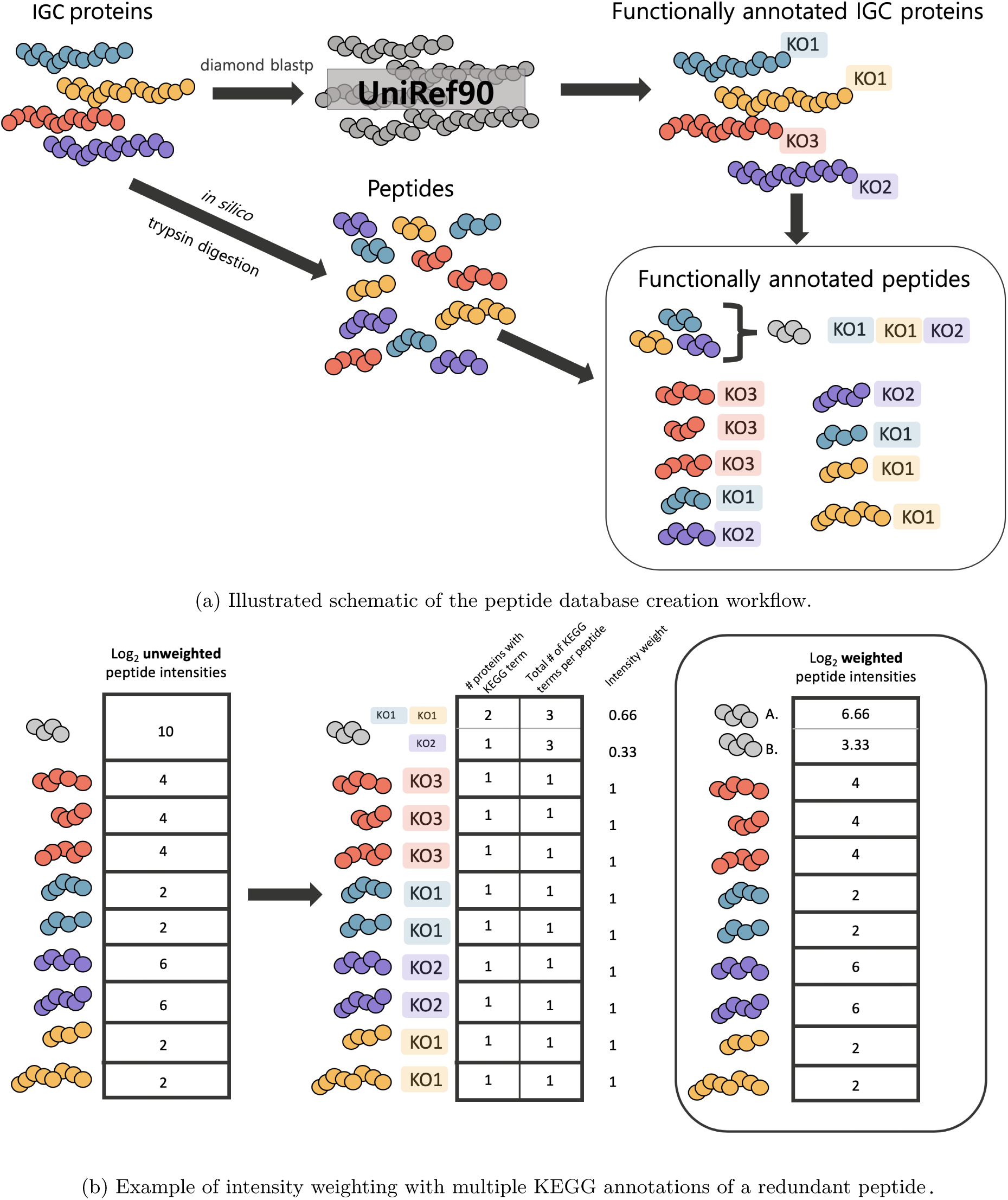
We first used diamond blastp to align IGC proteins to the UniRef90 gene cluster database for annotation of IGC proteins with KEGG terms. In parallel, we completed an *in silico* trypsin digestion of IGC proteins into tryptic peptides that then inherited KEGG annotations from their parent proteins. Notably, redundant peptides inherited all possible KEGG annotations. In this example, the redundant peptide (grey) can be found in three proteins. Thus, its intensity is weighted by our confidence in the peptide’s KEGG annotation. The redundant peptide is part of two proteins annotated with KO1 and one protein annotated with KO2, therefore we weight the peptide’s intensity for KO1 and KO2 by 0.66 (2/3) and 0.33 (1/3) respectively. The weighted intensities are then be used in our modified GSVA pipeline where the weighted intensities are associated to the appropriate peptide gene set.

After *in silico* trypsin digestion, our IGC database of 9,878,647 proteins consisted of 603,457,781 total peptides and 414,419,478 unique peptide sequences. For computational speed, we reduced this database size to peptides frequently matched in human gut microbiome studies to 469,393 unique peptides that were identified from 500 in house, raw MS/MSfiles. Of these unique peptides, 224,836 (47.9%) have KEGG annotation. Conversely, the IGC protein database has 2,109,127 (21.4%) proteins with KEGG annotation. The reduced database is made available as File S1.

### “Peptide”- and “protein”-set variation analysis

We adapted the GSVA method (Hänzelmann et al., 2013) for use with peptide intensity levels in metaproteomic experiments rather than gene expression estimation from RNA sequencing or microarray experiments. Peptide intensities were corrected by sample-specific size factors to normalize inter-sample variability. We calculated size factors using the `DESeq2` R package (Love et al., 2014) which consisted of median ratios of peptide intensities to the geometric mean of each peptide in the entire experiment. Peptide intensities were then divided by their corresponding sample-specific size factor. We removed peptides with intensities missing in over half the samples in each tested condition as a preprocessing filtering step for missing data. We then created “peptide gene sets” for our adapted GSVA analysis consisting of peptides annotated in each KEGG pathway for a total of 229 peptide gene sets. For our protein level analysis, we also created protein gene sets from protein groups annotated in each KEGG pathway. Peptides with multiple KEGG annotation terms were included in all appropriate gene sets by intensity weighting while considering the frequency of KEGG term annotation as explained in Equation 1 and Figure 1b:

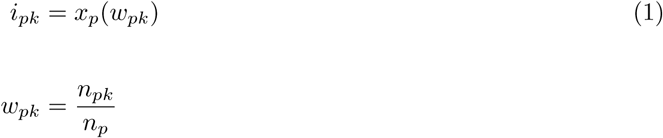

where *i* = adjusted intensity, *x* = measured peptide intensity, *w* = weight adjustment, *p* = peptide, *k* = KEGG term, *n* = number of KEGG terms.

We completed a peptide gene set variation analysis using the GSVA R package (Hänzelmann et al., 2013). We used a Gaussian cumulative distribution function for kernel estimation of each peptide’s intensity. To reduce noise, we only considered peptide gene sets with a minimum size of 10 peptides. The weighted peptide intensities (*w*_*pk*_) were further transformed by log_2_. Briefly, GSVA scores, a type of enrichment score, were calculated from ranking peptides in each set and calculating a Kolmogorov-Smirnov (KS) like random walk statistic from these ranked peptide sets. Significant differences in GSVA scores of peptide gene sets between tested conditions were identified using `lmFit()`, a least squares linear model, and `eBayes()`, empirical Bayes moderation, from the `limma` R package (Phipson et al., 2016) while considering multiple hypothesis testing by adjusting *p*-values using the Benjamini–Hochberg procedure (Benjamini and Hochberg, 1995).

We compared the results of the peptide gene set variation analysis workflow to that of a similar workflow centered on protein level intensity data. The methods remained the same, except we used protein group label-free quantitation (LFQ) intensity values provided by the MetaLab workflow. The protein LFQ values were also normalized by log_2_ transformations and then subjected to the same workflow as the peptide intensities including weighting of all protein group intensities with multiple KEGG terms.

### R shiny app construction

We created pepFunk, an R shiny application for our peptide-centric functional enrichment workflow. pepFunk was written in the R programming language v3.4.4 (R Core Team, 2019) and is dependant on the R packages: `shiny` (Chang et al., 2019), `shinydashboard` (Chang and Borges Ribeiro, 2018), `shinyWidgets` (Perrier et al., 2019), `DT` (Xie et al., 2019) for application building, `rhandsontable` (Owen, 2018), `reshape2` (Wickham, 2007), `tidyverse`, `plyr` (Wickham, 2011) for data manipulation, `colourpicker` (Attali, 2017), `tidyverse` (Wickham, 2017) and `plotly` (Sievert, 2018) for custom plotting, `DESeq2` (Love et al., 2014), `GSVA` (Hänzelmann et al., 2013) and `limma` (Ritchie et al., 2015) for data analysis, `ggdendro` (de Vries and Ripley, 2016) and `dendextend` (Galili, 2015) for dendrogram plotting, and `LaCroixColoR` (Bjork, 2019) for colour palettes. Dataset 1 is provided as sample data within the app, and was also deposited to the Pro-teomeXchange Consortium via the PRIDE (Perez-Riverol et al., 2019) partner repository with the dataset identifier PXD016388. Our application is hosted at https://shiny.imetalab.ca/pepFunk/ with source code available at https://github.com/northomics/pepFunk.

### Datasets

#### Dataset 1: Fecal microbiome treated with an HDAC inhibitor

We adopted an *ex vivo* microbiome assay, termed RapidAIM (rapid assay of individual’s microbiome) (Li et al., 2019) to assess the direct effects of the histone deacetylace (HDAC) inhibitor suberoulanilide hydroxamic acid (SAHA) on a human microbiome. Briefly, in a RapidAIM assay, a human gut microbiome (fecal) sample was cultured for 24 hours in anaerobic conditions in control conditions with dimethyl sulfoxide (DMSO) and treatment conditions with a low (0.125mg/ml), or high concentration of SAHA (0.25mg/ml). Proteins were digested with trypsin (Worthington Biochemical Corp., Lakewood, NJ). The digest was then desalted and analyzed using an Orbitrap Q-Exactive mass spectrometer as described previously (Zhang et al., 2018a). Spectra search and peptide quantitation were completed using MetaLab v1.1.1 (Cheng et al., 2017) and a database search of the IGC. The mass spectrometry proteomics data have been deposited to the ProteomeXchange Consortium via the PRIDE (Perez-Riverol et al., 2018) partner repository with the dataset identifier PXD016388. Peptide and protein group output files were used for the analyses. Of the total detected peptides, 79.5% were found in our core peptide database and 52.0% had at least one associated KEGG term (Figure S1a).

#### Dataset 2: Fecal microbiome treated with metformin

A human fecal sample (microbiome) stored at -80° C was thawed quickly at 37° C and cultured using RapidAIM (Li et al., 2019) for 24 hours with 10mM metformin (MTFM) or DMSO as the control. Control and treatment samples were cultured in five replicates. Cultured microbiome samples were subjected to protein extraction and tryptic digestion, and samples were analyzed using an Orbitrap Q-Exactive and a 90 min gradient as described previously (Li et al., 2019). Three technical replicates were run on the Q-Exactive for a single sample (MTFM_3). Spectra search and peptide quantitation was completed using MetaLab v1.2.0 (Cheng et al., 2017) using a database search of the IGC. The mass spectrometry proteomics data have been deposited to the ProteomeXchange Consortium via the PRIDE (Perez-Riverol et al., 2018) partner repository with the dataset identifier PXD016427. The median values of both peptide and protein level intensities were used for the analyses for the three technical replicates (MTFM_3). Of the total detected peptides, 84.4% were found in our core peptide database and 62.4% had at least one associated KEGG term (Figure S1b).

## Results

### Both peptide and protein level PCA can distinguish treatment groups

We first compared the ability principal component analysis (PCA) from intensity values at the protein- and peptide-levels distinguish treatment groups. To do so, we applied both protein and peptide-centric metaproteomic workflows to two fecal microbiome datasets: Dataset 1, a fecal microbiome treated with two concentrations of SAHA and Dataset 2, a fecal microbiome treated with metformin. PCA using both peptide and protein group intensities is able to separate the high concentration of SAHA from the control DMSO treatment of Dataset 1 (Figures 2a, 2b). Clustering of treatments is similar using peptide and protein level data analyses. In Dataset 2, PCA can also very clearly distinguish between a DMSO treated microbiome from one treated with metformin using both peptide and protein group intensities (Figures 2c, 2d). The control DMSO treatment samples, however, cluster more tightly when using peptide intensities.

**Figure 2:**
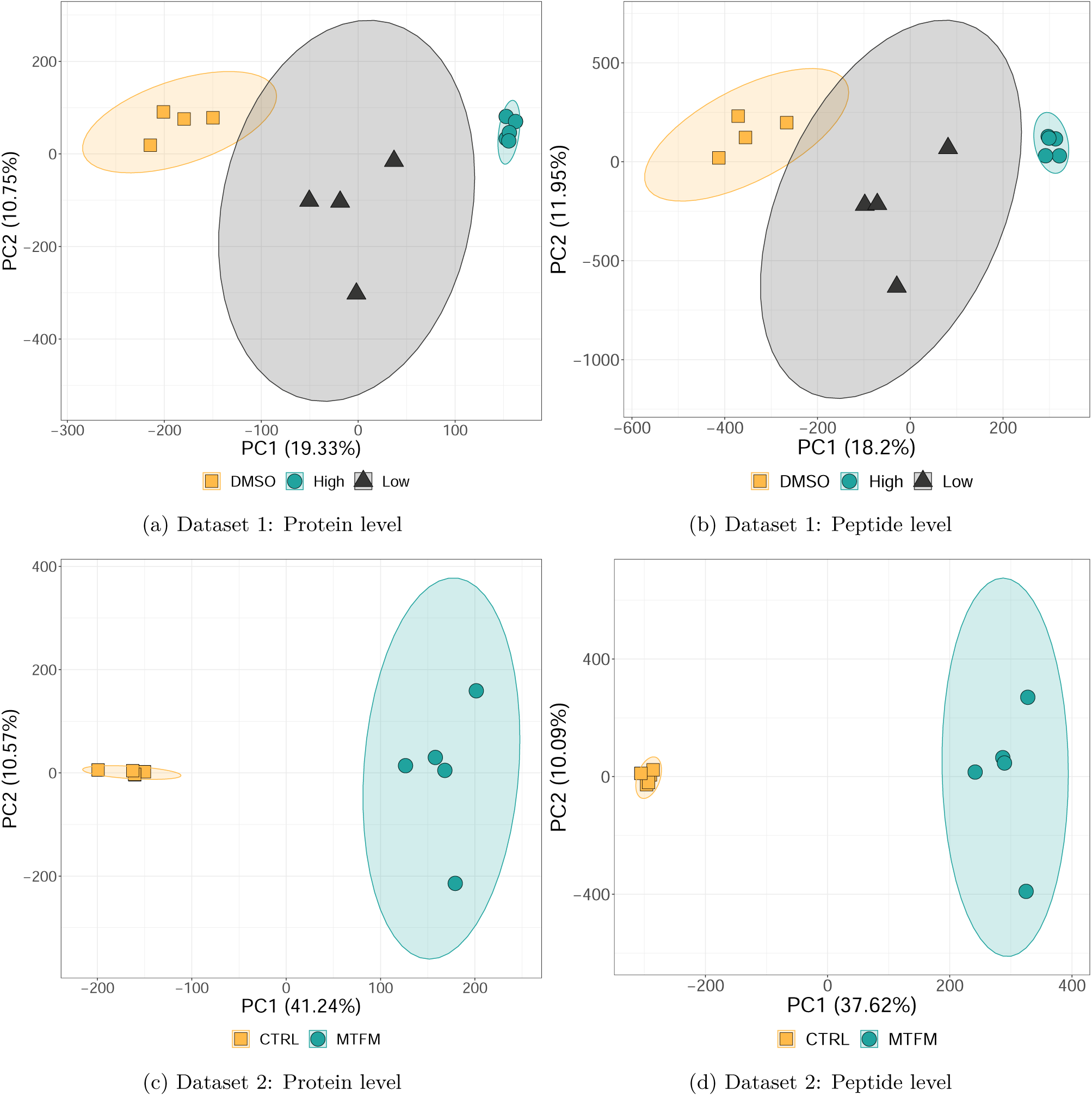
PCA biplots of Principal components (PC) 1 and 2 produced from both protein- and peptide-centric metaproteomic workflows of Datasets 1 and 2.

### KEGG functional enrichment of metaproteomic data

We compared protein- and peptide-centric workflows for KEGG pathway enrichment using a GSVA framework. Peptide spectra matches (PSMs) for both workflows were identified through a database search of the IGC peptide database using MetaLab (Cheng et al., 2017). Our protein level analysis considered protein groups that were identified by sequence similarity using MetaLab (Cheng et al., 2017) and KEGG annotation for all proteins in each protein group were considered for functional enrichment.

We computed GSVA scores for all samples in each dataset. To test if GSVA scores followed the same trend in both workflows, we completed a correlation analysis of the median GSVA scores of pathways found to be significantly enriched at either a peptide or protein level analysis. We found linear agreement of GSVA scores between using protein and peptide level data sources with Pearson’s correlation coefficients of 0.76 and 0.85 for Datasets 1 and 2 respectively (Figure 3).

**Figure 3:**
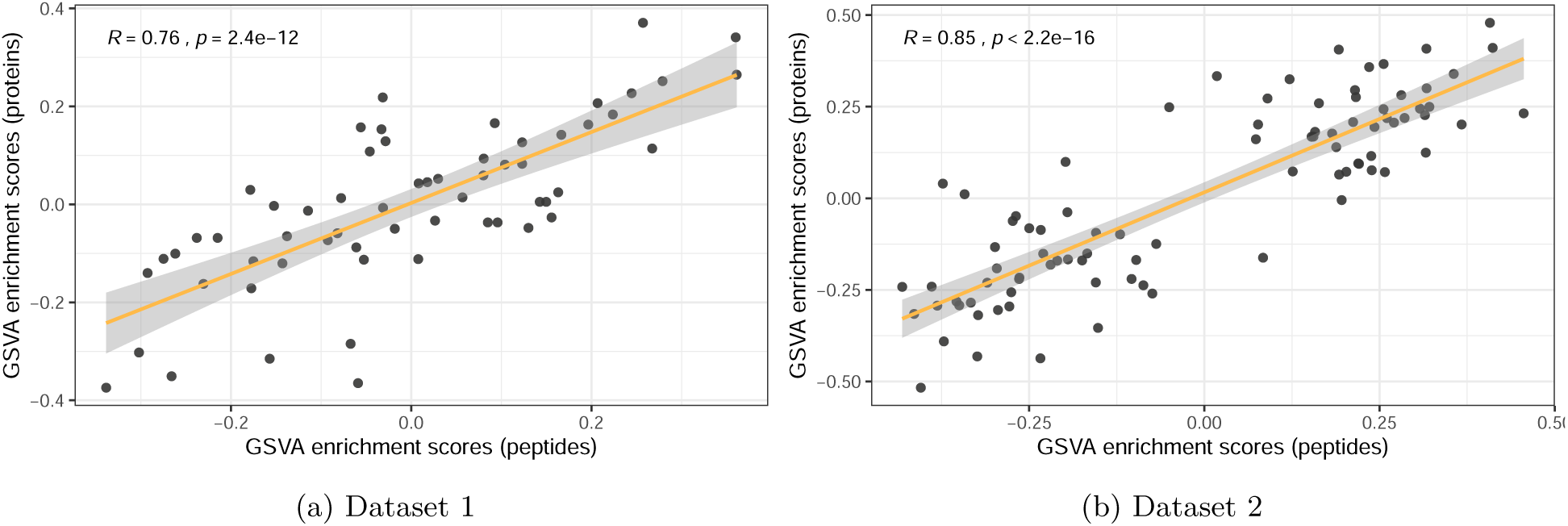
Correlation of significant peptide-centric and protein-centric GSVA scores. Median GSVA scores at the condition level were used for the analysis. A linear regression line is plotted in yellow with a grey ribbon representing a 95% confidence interval.

Using Dataset 1, we were able to complete GSVA on 70 and 87 protein and peptide gene sets respectively. After protein and peptide gene set GSVA score ranking, a linear model in combination with an Empirical Bayes approach, was used to identify differentially enriched KEGG pathways in each of the treatment conditions (high and low SAHA) compared to a control. Using the protein-centric workflow on Dataset 1, we identified two consistent and significantly enriched KEGG pathways when comparing control DMSO treatment to both concentrations of SAHA (selenocompound metabolism [PATH:KO00450] and biosynthesis of ansamycins [PATH:KO01051]; Figure 4). By only comparing control DMSO conditions to high concentrations of SAHA treated microbiomes, we identified four additional enriched KEGG pathways.

**Figure 4:**
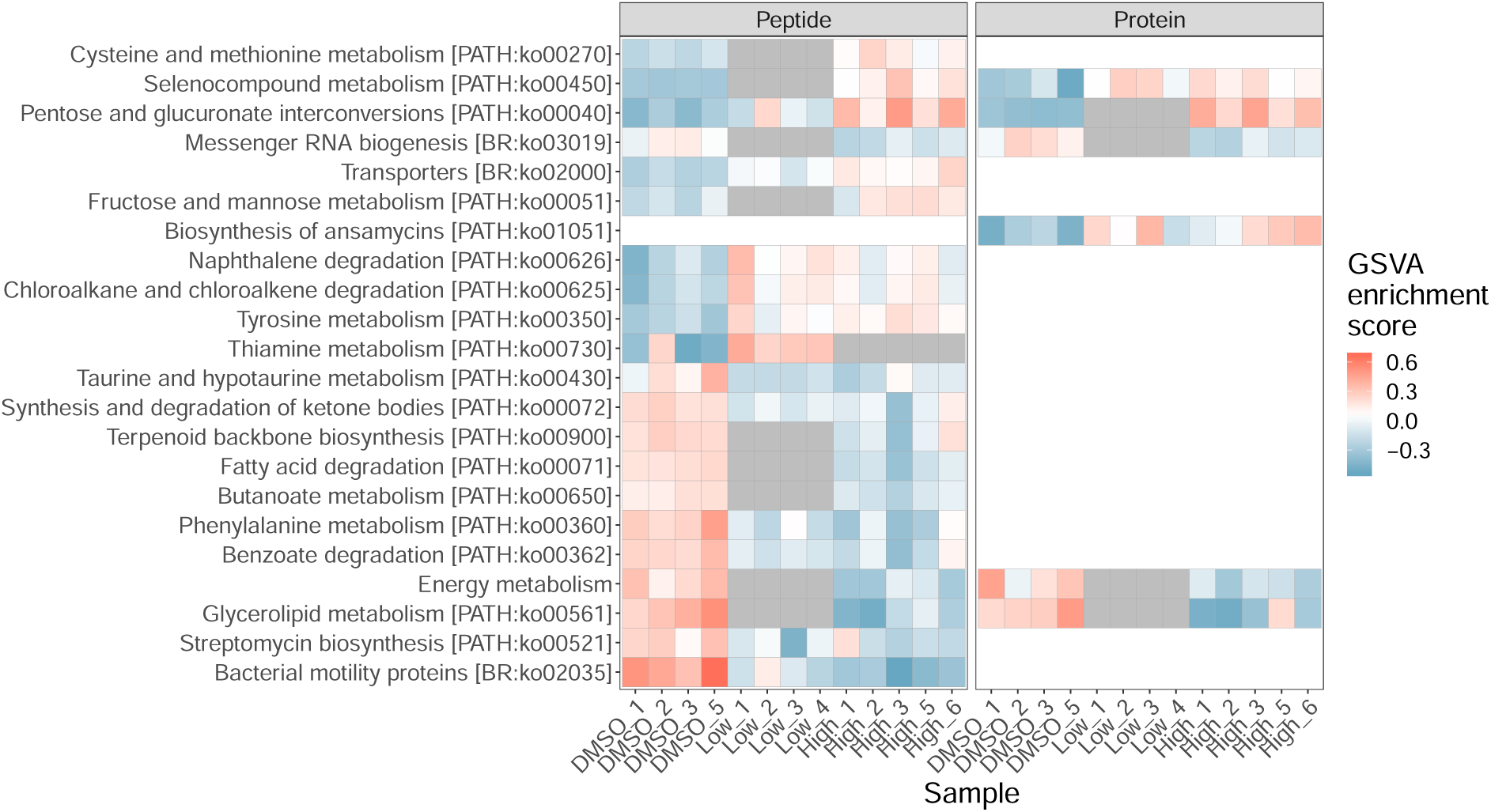
Heatmap visualizing GSVA scores of Dataset 1. Peptide centric workflow is presented on the left, and protein-centric on the right. A high score is visualized in coral and a low score in blue. A pathway is coloured in gray if the pathway is enriched in one condition, but not the other (BH adjusted *p* > 0.05). If the pathway is enriched in the peptide level analysis but not the protein level analysis, or vice versa, the pathway is not coloured in and is represented completely in white.

The peptide level analysis led us to identify twelve KEGG pathways as significantly enriched in samples treated with either low or high concentrations of SAHA (Figure 4). Thiamine metabolism was the sole pathway that was only significantly enriched in samples treated with a low concentration of SAHA, while nine other KEGG pathways such as glycerolipid metabolism and selenocompound metabolism were significantly altered after treatment with high levels of SAHA. Five of the same pathways were identified as significant by both peptide- and protein-centric approaches, however, the peptide-centric approach was able to identify more significantly enriched KEGG pathways than the protein-centric method (Figure S2). Notably, significantly enriched KEGG pathways identified by the peptide-centric workflow were enriched in the same direction in the protein-centric workflow in both datasets (Figure 4).

There was adequate detection of protein-groups for GSVA analysis on 67 KEGG pathway gene sets when considering the protein-centric analysis of Dataset 2. We identified 30 significant differentially enriched KEGG pathways in fecal microbiomes cultured with meformin. Conversely, we were able to complete GSVA on 94 gene sets using the peptide-centric approach, of which 47 were significantly enriched. Of the significantly enriched KEGG pathways, 24 were identified by both peptide- and protein-centric approaches (Figure S2c).

### R shiny app

We created a web-based peptide-centric workflow made available as a companion tool to MetaLab (Cheng et al., 2017) and iMetaLab (Liao et al., 2018). Our application, pepFunk, accepts input files as MaxQuant `peptide.txt` files or user formatted files that includes peptide sequence and intensity values. The app performs the entire workflow and allows for the user to visualize data as a PCA biplot and GSVA score heatplots. Users can also download analysed data to create their own customized figures. The app is available at https://shiny.imetalab.ca/pepFunk with source code found at https://github.com/northomics/pepFunk. Dataset 1 has been provided as sample data.

## Discussion

Functional analysis of metaproteomic data can be challenging. Database choice can have effects on the quality of results (Tanca et al., 2016), redundant peptides can lead to ambiguously identified proteins (Ning et al., 2016) and current methods of protein level analysis can lead to a loss of information. Typically, protein-centric workflows are used and can be considered analogous to a transcriptomic workflow where sequenced cDNA reads are mapped to genomic locations and analyses are completed on estimated transcript expression values. Recently, metatranscriptomics has moved towards functional annotation at cDNA read level which does not necessitate assembly or read mapping to genomic locations (Ugarte et al., 2018). However, instead of enzymatic digestion as seen in metaproteomics, fragmentation of cDNA for metatranscriptomic sample preparation can be performed by physical methods, such as sonication (Marine et al., 2011). The randomness of sonication typically leads to cDNA reads that map to unique locations in reference genomes, increasing the confidence of functional assignment.

In metaproteomics, the Protein Inference Problem, describing the challenges of peptide-to-protein assignment, can be even more difficult when considering the complexity of the microbiome. Because metaproteomic analyses can identify more proteins that share the same peptide sequences through the inclusion of multiple microbial strains and species, protein group level analyses have been used to analyze proteins clustered into groups by sequence similarity. However, assigning peptides to protein-groups leads to data loss where researchers can lose statistical power and potentially important functional information of their microbial community samples (*e.g.* Figures 4,5). To combat the issues that can arise from protein group pipelines we have created a peptide-centric workflow. By analysing metaproteomic data at the peptide level, we are able to identify similar enriched KEGG pathways as analysis at the protein group level. Furthermore, we can identify more enriched KEGG pathways at the peptide level compared to the protein level because we retain more information (*e.g.* 12 vs 6 in Dataset 1 and 47 vs 31 in Dataset 2; Figures 4, 5, S2). Our peptide-centric workflow is unique as it uses a weighted intensity for functional assignment that is proportional to our confidence in annotated KEGG terms. In addition, our database is small and reduces computational resources required for a full database search.

**Figure 5:**
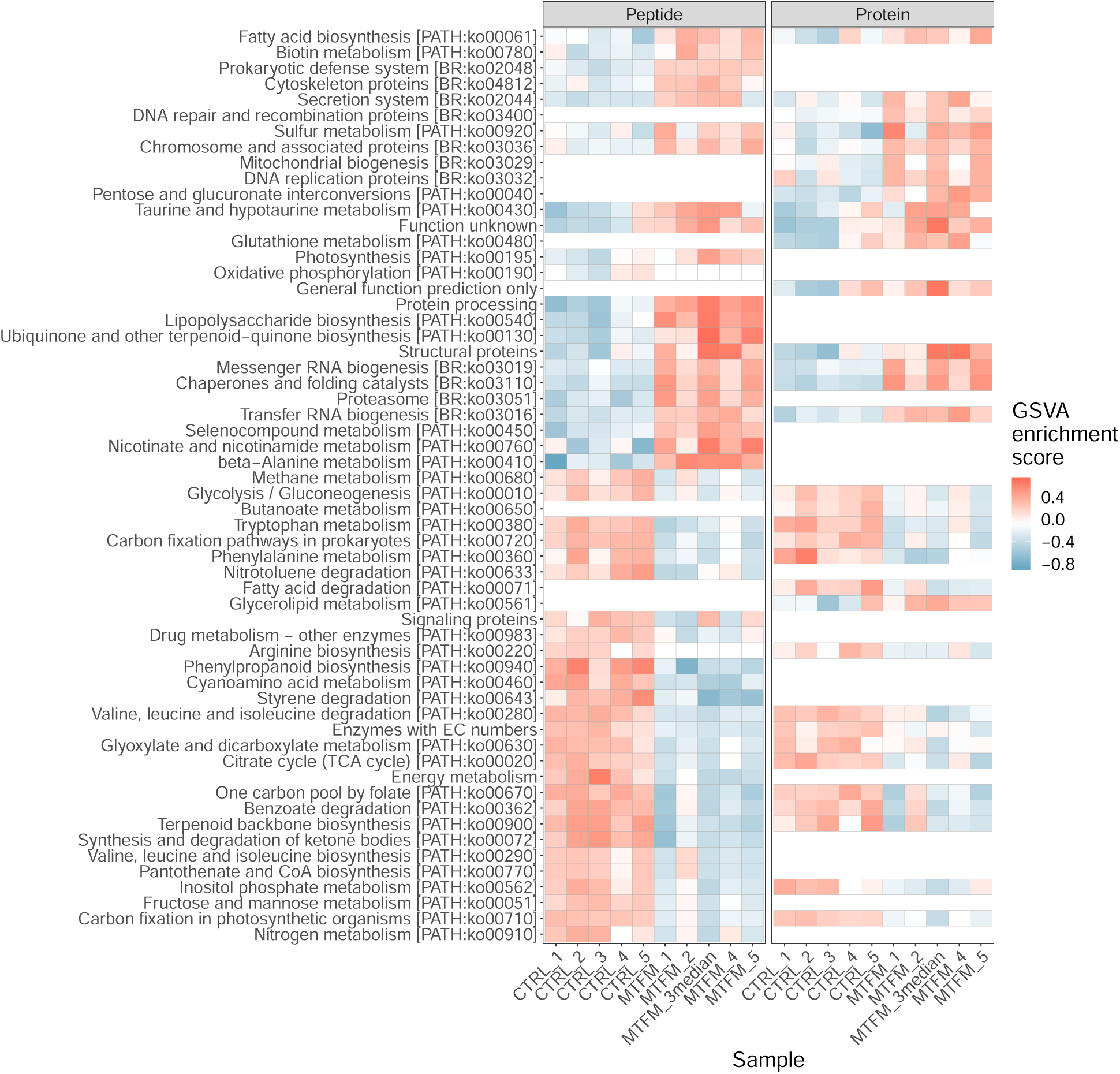
Heatmap visualizing GSVA scores of Dataset 2. Peptide centric workflow is presented on the left, and protein-centric on the right. A high score is visualized in coral and a low score in blue. If the pathway is enriched in the peptide level analysis but not the protein level analysis, or vice versa, the pathway is not coloured in and is represented completely in white.

To confirm the appropriateness of our approach, we looked at the biological relevance of the enriched KEGG pathways in our peptide-centric results. For Dataset 1, a microbiome treated with SAHA, an HDAC inhibitor, we looked at the cited functional roles of acetylation in bacteria. Acetylation is a reversible post-translational modification most well known for being essential to gene regulation. Acetylation can epigenetically alter expression by reducing the interaction between histones and DNA making DNA more accessible to transcriptional machinery (Sterner and Berger, 2000). However, although HDAC inhibitors increase acetylation, SAHA and other inhibitors are also known to repress transcription (Greer et al., 2015) thus reduced expression of peptides associated to mRNA biogenesis [BR:KO03019] is expected (Figure 4).

Castaño-Cerezo et al. (2014) used *cobB* and *patZ* knockout mutants (a deacetylace and an acetyltrans-ferase) in *Escherichia coli* to study how acetylation can affect the bacterial species. We expected to see similar functional results in our study to that by Castaño-Cerezo et al. (2014), particularly the results of the *cobB* knockout *E. coli*. Castaño-Cerezo et al. (2014) identified that 64% of acetylated proteins were associated to metabolism. Similarly, after SAHA treatment, we identified alterations in expression to many pathways associated to metabolism, such as increases in cysteine and methionine [PATH:KO00270] and selanocompound [PATH:KO00450] metabolism, and decreases in butanotate [PATH:KO00650], phenylalanine [PATH:KO00360], and benzoate [PATH:KO00362] metabolism (Figure 4). Additionally, there is evidence to show that acetate metabolism itself is affected by acetylation through acetyl-CoA generation by acetyl-CoA synthase (ACS) (Castaño-Cerezo et al., 2014). For example, CobB was shown to preferentially deacetylate ACS, increasing its activity. The *cobB* mutant displayed reduced ACS activity, thus suggesting a reduction in acetyl-CoA generation. We identified a reduction in both fatty acid degradation [PATH:KO00071] and butanoate metabolism [PATH:KO00362] pathways (Figure 4), both of which result in acetyl-CoA. Reduced acetyl-CoA by HDAC inhibition may also be decreasing acetate metabolism. However, we identified a reduction in cell motility when gut microbes were treated with SAHA, the opposite finding of Castaño-Cerezo et al. (2014).

Metformin, a drug widely used in the treatment of type II diabetes, has previously been shown to alter gut microbiome taxonomic composition and functionality (De La Cuesta-Zuluaga et al., 2017; Ma et al., 2018; Li et al., 2019). In our study, we identified metformin-induced alterations to the same pathways as other studies, for example a decrease in fatty acid biosynthesis [PATH:KO00061] (Li et al., 2019), decrease in nitrogen metabolism [PATH:KO00910], increase in oxidative phosphorylation [PATH:KO00190], enrichment in lipopolysaccharide biosynthesis [PATH:KO00540] (Ma et al., 2018), and an increase in tRNA biosynthesis [BR:KO03016]. Metformin has also been shown to reduce folate metabolism (Cabreiro et al., 2013), thus lower “one carbon pool by folate” GSVA scores in metformin treated samples are expected. However, we identified a significant reduction in peptide intensity associated to glycolysis/gluconeogenesis in our metformin treated samples, the inverse of the findings by Li et al. (2019). This finding may be due to an abundance of proteolytic bacteria in this sample, or to the comparison between our human samples to the mouse samples from (Li et al., 2019).

Currently, the vast majority of gut microbiome studies focus on using genomic sequencing to identify microorganisms. This type of meta”-omics”, is useful at identifying the composition of a microbial community or its corresponding functional potential using a shotgun metagenomics approach (Halfvarson et al., 2017). However, metagenomics cannot identify if genes are expressed and functionally active. Identifying the functionality of a microbiome sample is essential because multiple taxa can have redundant functions, and it is possible that a microbiome persists functionally even when taxa composition is altered (Blakeley-Ruiz et al., 2019). As such, the emerging field of metaproteomics instead offers a functional snapshot of microbiome by identifying and quantifying translated proteins. Recently, multi-omic studies have shown that taxonomic variation, identified through metagenomics, may not always be associated with overall metaproteomic-identified functional changes in microbiome studies (Blakeley-Ruiz et al., 2019; Mikan et al., 2019). For example, Blakeley-Ruiz et al. (2019) identified compositional taxonomic changes in the microbiomes of patients with inflammatory bowel disease (IBD) yet persistent metabolic functionality both within and between patients. While it is accepted that the taxonomic composition and abundance in gut microbiomes are variable between individuals (Yatsunenko et al., 2012), it is possible that redundancy in microbe functions result in an invariable metabolic landscape between individuals. Thus if taxonomic changes do not always lead to functional shifts, metaproteomic studies should also consider taxa-independent functional analyses of samples, a focus of our peptide-centric workflow. Additionally, peptide-centric taxonomic analysis of metaproteomic data is already implemented by MetaLab (Cheng et al., 2017), therefore our peptide functional analysis completes the data analysis workflow and demonstrates the merit of working towards completely peptide-centric metaproteomic data analysis in gut microbiome studies.

GSVA, our functional analysis method of choice, can be used in a condition-independent manner in addition to comparing between control and treatment samples (Hänzelmann et al., 2013). A condition-independent type of analysis is useful for researchers using functional pathways for exploratory analysis or for observing pathways that may be highly or lowly expressed in any given sample. Hänzelmann et al. (2013) also demonstrated that GSVA analysis has a higher degree of sensitivity than other gene set enrichment techniques, such as ssGSEA and PLAGE, while simultaneously maintaining a low type-I error rate of approximately 0.05. While differential protein expression analysis can be used to study metaproteomic data (Hamann et al., 2016), differential expression analysis can only identify differences of expression of individual proteins. Our implementation of GSVA gives sample- or experiment-wide interpretable results associated to KEGG pathways with a functional database specific to human gut microbiome studies. Currently, there also exists PSEA-Quant, which is a protein-centric gene set enrichment tool capable of performing both condition dependant and independent analyses, that uses protein gene set enrichment analysis, a method similar to our study (Lavallée-Adam et al., 2014; Lavallée-Adam et al., 2015; Lavallée-Adam and Yates III, 2016). However PSEA-Quant, uses protein level data from proteomic studies, and is not specific to human gut microbiome experiments.

As the field of metaproteomics grows, so does the need for accurate, fast, and user-friendly tools for data analysis. Current protein focused functional enrichment workflows struggle with data loss stemming from assigning redundant peptides to proteins or protein groups. To combat this challenge, we created and implemented pepFunk, a peptide-centric functional enrichment workflow and accompanying R shiny tool for accurate and customizable data analysis. The current version includes a custom KEGG to human gut microbiome peptide functional database, but more experienced users can use their own annotated peptide database. As Ning et al. (2016) proposed, we have developed a workflow that directly analyses peptide intensities and is able to identify enriched KEGG pathways while maintaining the statistical validity of a protein-centric approach.

## Additional files

File S1: Annotated core peptide database.

File S2: Supplemental figures (follows manuscript)

---

**Figure S1:**
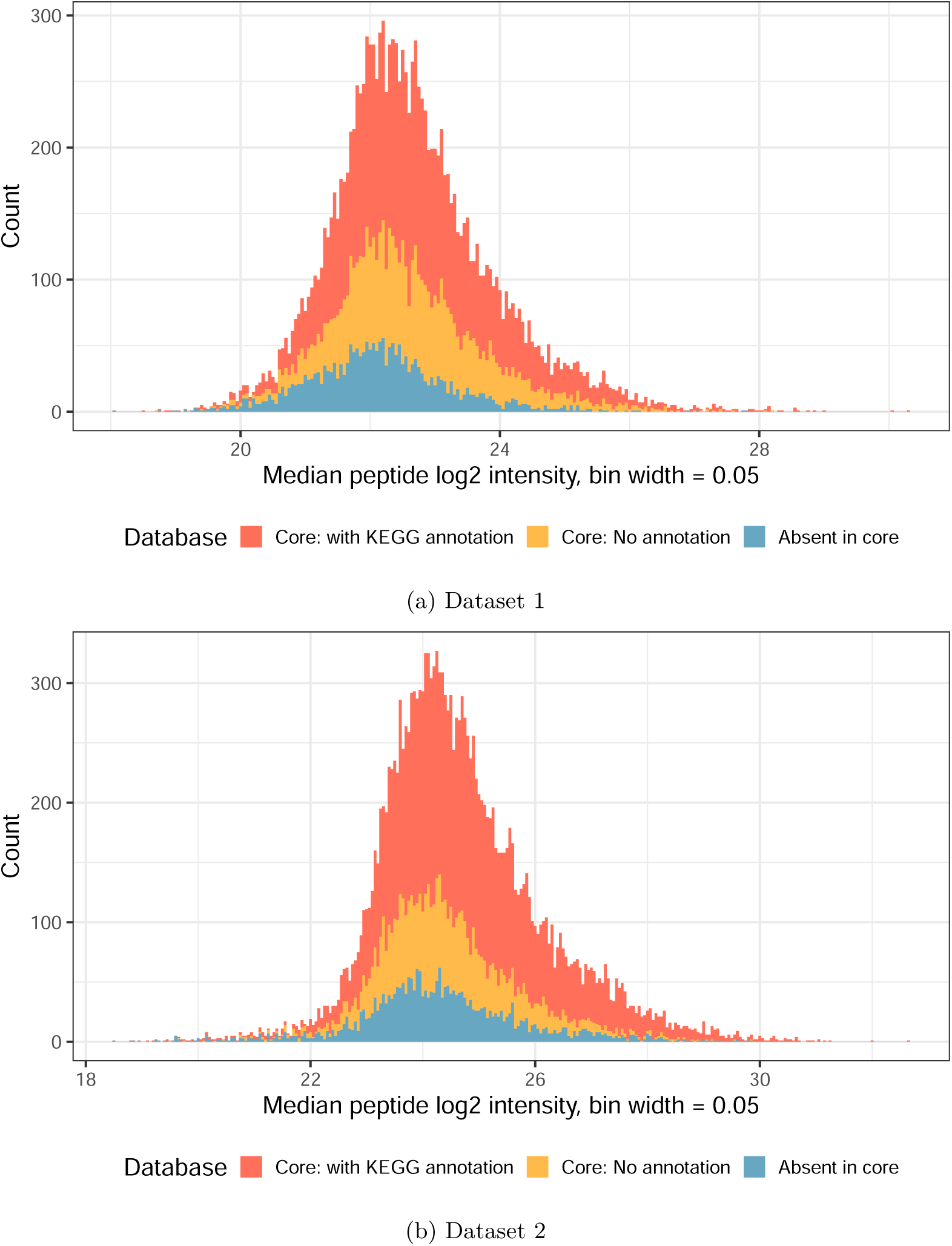
Frequency distribution of median log_2_ peptide intensity values found in core peptide database. Missing data (intensity values of 0) were removed.

**Figure S2:**
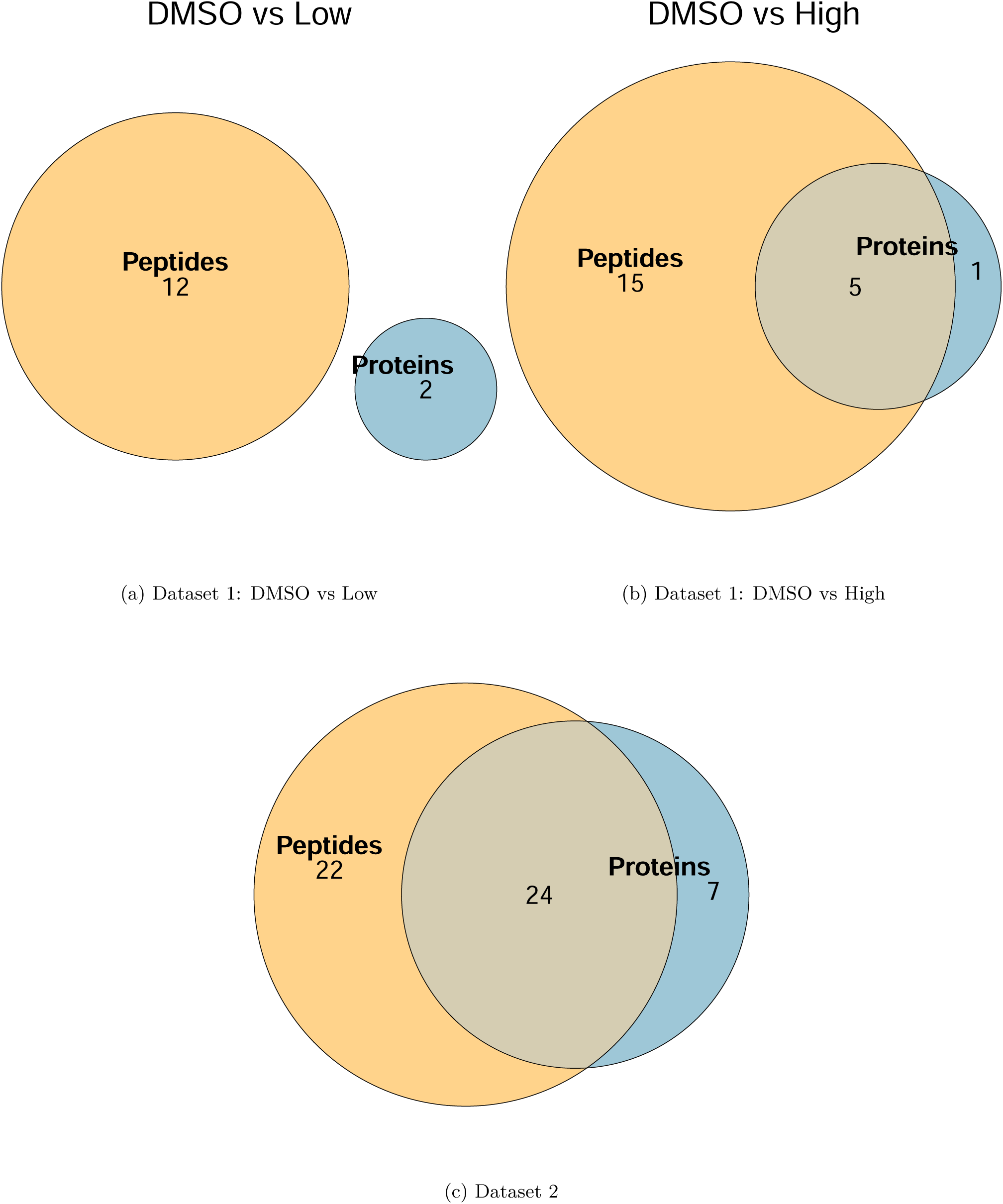
Venn diagrams depicting the overlap in enriched KEGG pathways identified using a GSVA framework and peptide vs protein level data.

